# Murine bone properties and their relationship to gait during growth

**DOI:** 10.1101/465948

**Authors:** Hyunggwi Song, John D. Polk, Mariana E. Kersh

## Abstract

Allometric relationships have been queried over orders of mammals to understand how bone accommodates the mechanical demands associated with increasing mass. However, less attention has been given to the scaling of bone within a single lifetime. We aimed to determine if bone morphology and apparent density is related to (1) bending and compressive strength, and (2) gait dynamics. Longitudinal in vivo computed tomography and gait data were collected from female rats (n=5, age 8 - 20 weeks). Cross sectional properties and apparent density were measured at the diaphysis, distal, and proximal regions of the tibia and scaling exponents were calculated. Finite-element models were used to simulate four-point bending and axial compression using time-specific ground reaction forces (GRF) to calculate the mean strain energy density (SED) at the midshaft. Second moment of area at the diaphysis followed strain similarity based allometry, while bone area was positively allometric. The average SED at the diaphysis decreased, especially after the age of 10 weeks (R^2^=0.99), while it increased in compression (R^2^=0.96). The apparent density in all regions initially increased and converged by 11 weeks of age and this was correlated with changes in joint angle. The scaling analyses implies that rodent tibia is (re)modeled in order to sustain bending at the midshaft during growth. The finite element results and relatively constant density after 10 weeks of age indicate that structural parameters may be the primary determinant of bone strength in the growing rodent tibia. The correlations between bone properties and joint angles imply that the changes in posture may affect bone growth in specific regions.

## Introduction

The capacity for bone to accommodate a wide variety of sizes in mammals is a remarkable result of a mechanobiologically driven process. The material and structural properties of bone are organized during growth – when mass and size increase significantly – in order to sustain the loading cycles from everyday movement [1–3]. Mechanical stimuli on bone during these movements promote bone modeling and remodeling and are influenced by the loading direction determined in part by overall body posture [4, 5]. Understanding how bone accommodates the increases in mass associated with growth, presumably to minimize fracture risk, has been a longstanding subject of interest.

Fracture risk depends on the magnitude and orientation of the forces experienced in the limbs, the structural integrity of each segment [1, 6], and the material composition of bone. Force magnitudes are related to body mass and speed [1, 2, 7], while Song *et al*. Page 2 of 13 bone structural integrity is reflected in cross-sectional properties (CSP) including bone area (BA) and second moment of area (I) [5, 8, 9]. Cortical bone area, rather than total area, has been related to a bone’s resistance to tensile and compressive loading, while second moment of area reflects resistance to bending loads. Mineral density is indicative of the Young’s modulus of bone, and in humans is used as a clinical measure of fracture risk [10]. Bone mineral density has been shown to increase during growth in many species of mammals including rabbits, mice, rats, and humans [11–14].

The relationship between bone cross-sectional properties, CSP, and body mass, m, has been shown to follow an exponential function characterized by the scaling exponent a such that CSP ma. Structures that scale under “isometric” conditions exhibit constant proportional changes in size with respect to mass, resulting in a scaling exponent of 0.67 for properties related to area (e.g. bone area) and 1.33 for properties related to bending (e.g. second moment of inertia). However, isometric scaling results in increases in stress since mass increases with (length)^3^, but cross-sectional area increases with (length)^2^ [15–17]. Structures with scaling exponents that differ from isometric scaling are referred to as following allometric scaling. Structures with positive allometric scaling exponents equal to 1 for area, and 1.83 for second moment of area, maintain a constant stress with increases in mass, and structures that exceed these constant strain exponents exhibit decreases in stress with increasing mass.

In his initial observations, Galileo proposed that animals scale with positive allometry, exponents greater than isometric values, thereby offering an explanation for why large animals have relatively thicker bones compared to small animals. Scaling analyses have since been used to investigate how long bone cross-sectional properties change with body mass among several Orders of mammals and birds in order to determine if fracture risk changes with increases in mass [1, 2, 15, 17–22]. Some studies have noted that isometric or negative allometry was conserved regardless of species [23, 24]. For example, primates, rodents and carnivorans have been shown to scale similarly when bone length and mass was accounted for [22]. The long bones of small terrestrial carnivores have also been shown to exhibit slight negative allometry, but there was a strong positive allometry in large mammals [2, 25]. Decreased bone length has been suggested as a mechanism for minimizing bending stress in larger mammals but not so in smaller mammals [18]. However, the long bones of small mammals have been shown to exhibit positive allometry with respect to length, but no cross-sectional properties were measured in this study [26]. Recently, a study of the pelvic limb in small avians found positive allometric scaling [19], and the same trend was found in quadrupedal terrestrial tetrapods [18, 23, 27]. There is continued debate over whether or not the bones of small mammals follow isometric or allometric scaling with respect to mass.

In addition, few longitudinal studies have been performed during growth to determine if bone (re)modeling occurs to minimize strain energy and therefore decrease fracture risk. Further, it is unknown if the same scaling principles observed across multiple Orders of differently-sized mammals applies ontogenetically within a single species or even among closely related species. Most of the scaling work has involved broad interspecific studies e.g. phylogenetic groups (Rodents, Felids, and Primates, respectively) [2, 28, 29], but there are relatively few ontogenetic studies of bone allometry. In addition, scaling analyses are limited by their assumption of constant density and do not account for the continuum mechanics of bone loading since measurements are limited to a given cross-section of bone.

The aims of this study were to (1) characterize changes in bone morphology and apparent density changes during growth within a single species, (2) determine if changes in bone morphology and density are related to changes in joint kinematics, and (3) identify if bone structure is optimized for bending or compressive loads during growth. Using rats as a model system, longitudinal in vivo micro-computed tomography (CT) scans were performed at seven time points to measure rodent tibia bone morphology, apparent density during early growth. In parallel, gait kinematics and kinetic data were collected. We calculated the mass scaling exponent of bone area and second moment of area using cross-sectional CT data at the proximal, distal, and diaphyseal regions of the tibia. These analyses were supplemented by finite-element analyses of the tibia under bending and compression to account for material properties and the distribution of load along the tibia.

## Methods

### 1. Animal care and training

Five healthy female Sprague-Dawley rats were used for this study beginning at the age of 7 weeks (average weight = 205.00 ± 13.97 g, Charles River Laboratories, Inc.). Data were collected weekly until 12 weeks of age, followed by data collection at 14 and 20 weeks of age. Rodents were group housed in a humidity and temperature controlled room on a 12:12 hour dark-light cycle. Food and water were provided ad libitum. Rodents were allowed to adapt to their new environment and walking platform for one week. All protocols were reviewed and approved by the Institutional Animal Care and Use Committee (IACUC) at University of Illinois at Urbana-Champaign. Experiments were performed at the Beckman Institute of Advanced Science and Technology.

### 2. Bone imaging and image analysis

In vivo micro-computed tomography data (isotropic resolution= 35.8 *µ*m, Inveon PET/SPECT/CT, Siemens, Munich, Germany) of the hind limbs were collected at each time point (Fig. 1A,B). Three hydroxyapatite (HA) phantoms were included in all scans (Model 092, CIRS) to convert Hounsfield Units to apparent HA density. The right hind limbs of the rats were segmented semi-automatically from the CT images using Amira 5.6 (Visage Imaging GmbH, Berlin, Germany). An initial threshold was used by visual inspection to separate the bone from soft tissue followed by manual corrections of the outer bone contour to create a mask. Within the mask, bone was identified using the Iterative Self-Organizing Data Analysis Technique (ISODATA) [30] implemented in Matlab (MATLAB R2016b, MathWorks, Natick MA).

**Figure 1.**
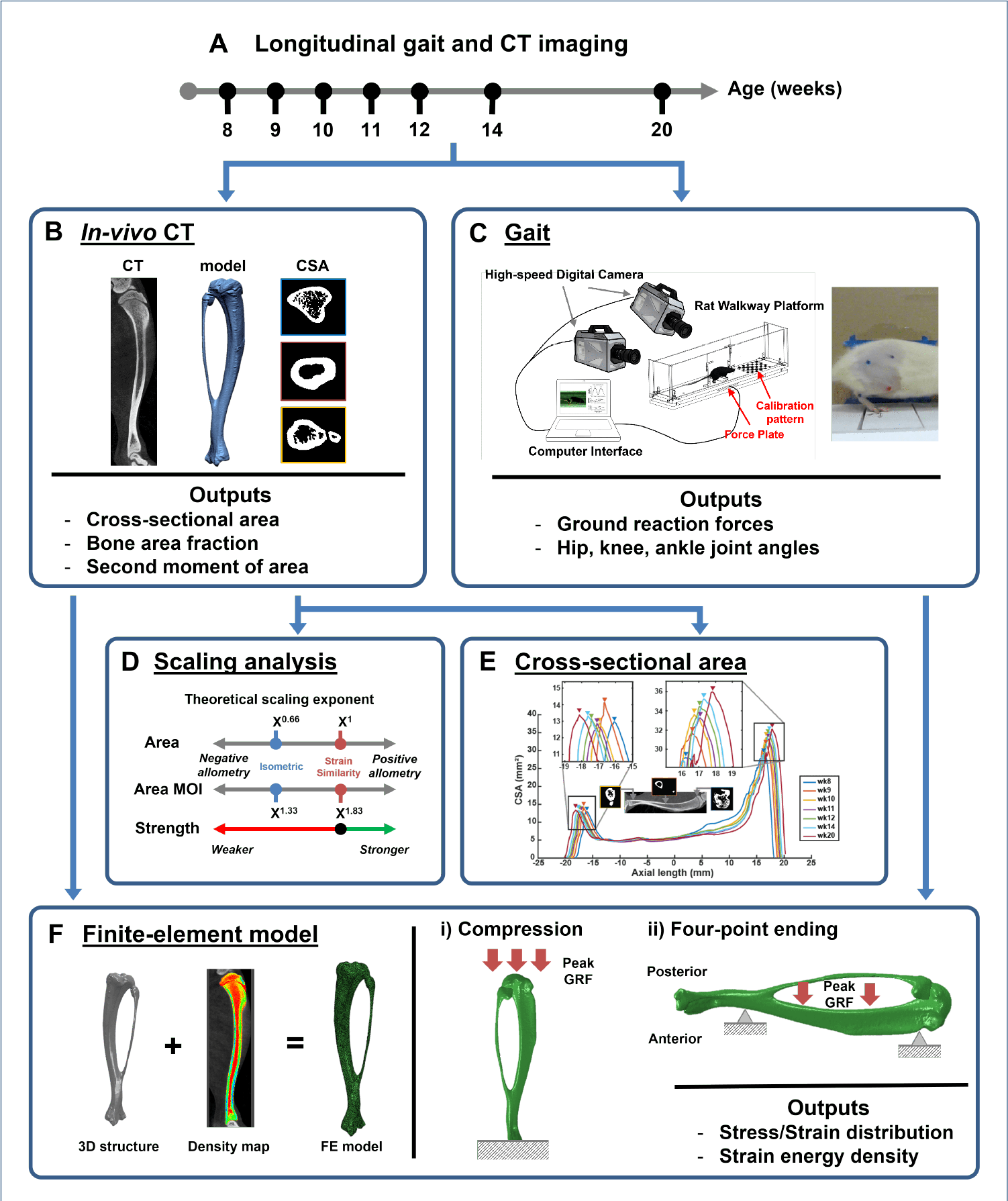
(A) Longitudinal musculoskeletal growth in the tibia was quantified between 8 and 20 weeks of age. (B) While under anesthesia, rats were imaged in vivo with microCT and the apparent density conversion was calculated. (C) Two images of the rat during gait were taken simultaneously by two high-speed digital cameras. Markers were defined on anatomical landmarks (iliac crest, hip, knee, and ankle). Joint kinematics during gait was obtained in 3D. The ground reaction forces were collected from the in-ground force plate. (D) Scaling analysis of bone area and second moment of area was performed. Theoretical scaling exponents of isometric growth and stress similarity based growth for each measurement were shown. (E) Cross-sectional properties and the apparent density were analyzed at proximal, midshaft, and distal regions of rat tibia. (F) Each element of 3D tibial structure was assigned an average Young’s modulus based on the local apparent density. For each model, compression loading and four-point bending were simulated with rat-specific GRFs at each time point to calculate the mean strain energy density at the midshaft.

To align the CT data on the same axis, two geometrical landmarks at the distal and proximal ends, were identified, and the CT image stack was resliced orthogonally along these reference points. For each bone, the midpoint of the total length of the bone was defined as the zero position in all samples. Our initial analyses of cross-sectional area revealed increases in CSA at the distal and proximal ends (Fig. 1E).

Therefore, we developed code to calculate the total transverse cross-sectional area, bone area (defined as the sum of cortical and trabecular area), apparent density, and bone area fraction (BA/TA) at the distal, proximal, and mid-slices. Apparent density of each slice was calculated as the average density of all segmented voxels. Second moment of area, which estimates the structure’s resistance to bending, along the bone was calculated with segmented bone area images using BoneJ (ImageJ, NIH) [31].

### 3. Gait analysis

Joint angles and ground reaction forces were measured during walking along a custom walkway with an isolated in-ground force sensor (Nano43, ATI Industrial Automation) (Fig. 1C). Prior to gait collection, rodents were anesthetized with isoflurane delivered by precision vaporizer after which the right hind limb and dorsum were shaved and five anatomical landmarks were marked with non-toxic permanent ink: the iliac crest, the greater trochanter of the femur, the lateral tibial tuberosity, the talocrural joint, and the distal and lateral aspect of the 5th metatarsal.

At each time point, rats were weighed and the kinematics of the right hind limb during walking was measured using two high-speed cameras. Each rat was placed on the walkway, and the joint kinematics and the ground reaction force data of the rat were recorded at the same frequency of 240 Hz. The positions of the surface markers were manually identified within each frame of the video data in Matlab. The three-dimensional positions of all surface markers were triangulated from both cameras using a calibration pattern included on the walkway [32]. The knee marker position was estimated in 3D by measuring the length of the femur and the tibia to minimize the effect of skin movement. The knee position was calculated by assuming that the estimated position of the knee marker would lie on the same plane defined by the hip, knee, and ankle markers as described in [33, 34]. Hip, knee, and ankle joint angles from heel strike to subsequent heel strike were calculated.

### 4. Finite element analysis of strain energy under bending and compressive loads

Finite element models were used to evaluate the strain energy within the mid-diaphysis during growth due to bending and compressive loads. The tibia of a representative rat was converted into finite-element model using an established computational pipeline [35]. Briefly, each segmented tibia was exported as triangulated surfaces and converted to solid models using a reverse engineering software (Geomagic). The solid models were meshed with quadratic tetrahedral elements (Abaqus, Simulia). Using a hydroxyapatite calibration phantom, CT hounsfield unit values were converted to apparent mineral density, and each element was assigned an average Young’s modulus based on the local apparent density following the relationships described in Cory et al. (2010) [36].

For each model, four-point bending and axial compression loading was simulated independently (Fig. 1F). We accounted for the increase in loading associated with growth by using rat-specific GRFs at each time point as the loading boundary condition. For compression, the peak ground reaction force at each time point was distributed over the tibial plateau while the distal end of the tibia was held fixed. For four-point bending, the peak ground reaction force at each time point was applied anteriorly at 35% and 65% of the axial bone length at the center of the posterior surface. A small region on the anterior surface at 20% and 80% of the axial bone length was held completely fixed. For each time point specific loading condition and geometry, the mean strain energy density (SED) within the cortical bone at the mid-shaft was calculated.

### 5. Data analysis

A repeated measures Analysis of Variance (ANOVA) was used to compare differences in peak ground reaction forces and joint angles over time. A fixed effects regression model was used to examine the correlation between gait parameters (peak forces and joint angles) and bone properties (cross-sectional area, density, BV/TV and BA/TA) during growth. This model held constant average effects of each rat. All ANOVA and regression analyses were performed with OriginPro 2018 (OriginLab, Northampton, MA, USA). All kinematic and kinetic data are shown as mean ± standard error (s.e.) and a Type I error rate *α*<0.05 was used for statistical comparisons.

Structural and mass data were transformed to log-log scales to allow for linear regressions between structural parameters and body mass. Bone area and second moment of area versus body mass were compared with isometric scaling exponents (0.67 for area and 1.33 for second moment of area) to investigate how biological characteristics change with size. Significant deviations from isometry were detected when the confidence limits associated with the linear fit excluded the isometric expectations.

Linear regressions between body mass and mean strain energy density within the cortical bone at the mid-shaft were used to assess increases or decreases in bone strength during growth. Correlation coefficients (r) over all time points were compared to those calculated using the last four time points.

## Results

From 8 to 20 weeks of age, the tibia bone length increased 11.7% with an initial rate of longitudinal growth of 1.01 ± 0.13 mm/week and decreased to 0.18 ± 0.01 mm/week by age 20 weeks. As expected, the proximal region of the tibia had the largest cross-sectional area (CSA) which increased from 32.0 ± 2.2 mm^2^ at 8 weeks of age to 36.8 ± 2.2 mm^2^ by 12 weeks (p<0.05) (Fig, 2A). Increase in proximal CSA (1.19 ± 0.49 mm^2^/week) was greatest before 11 weeks of age, after which the proximal CSA growth rate decreased to 0.14 ± 0.27 mm^2^/week. The diaphyseal and distal CSAs were constant between 8 and 20 weeks of age. However, bone area fraction (BA/TA) within the diaphysis and distal tibia increased. Distal BA/TA increased until 10 weeks and then exhibited a small, but not significant decrease beyond ten weeks of age (Fig. 2B). Diaphyseal BA/TA increased 11.5% peaking at 14 weeks (p<0.05) after which there was also no change. Proximal BA/TA was more variable during growth: it increased 7.9% between 8 and 12 weeks of age, decreased by 5.1% at 14 weeks but again increased 7.7% at 20 weeks of age. By 20 weeks of age, the bone area fraction for all three regions converged to 0.69 ± 0.01. Finally, density in all three regions of the tibia increased 38.6, 31.5, and 23.1% from 8 weeks of age until 11 weeks for the proximal, diaphyseal, and distal regions respectively (p<0.05, Fig. 2C).

**Figure 2.**
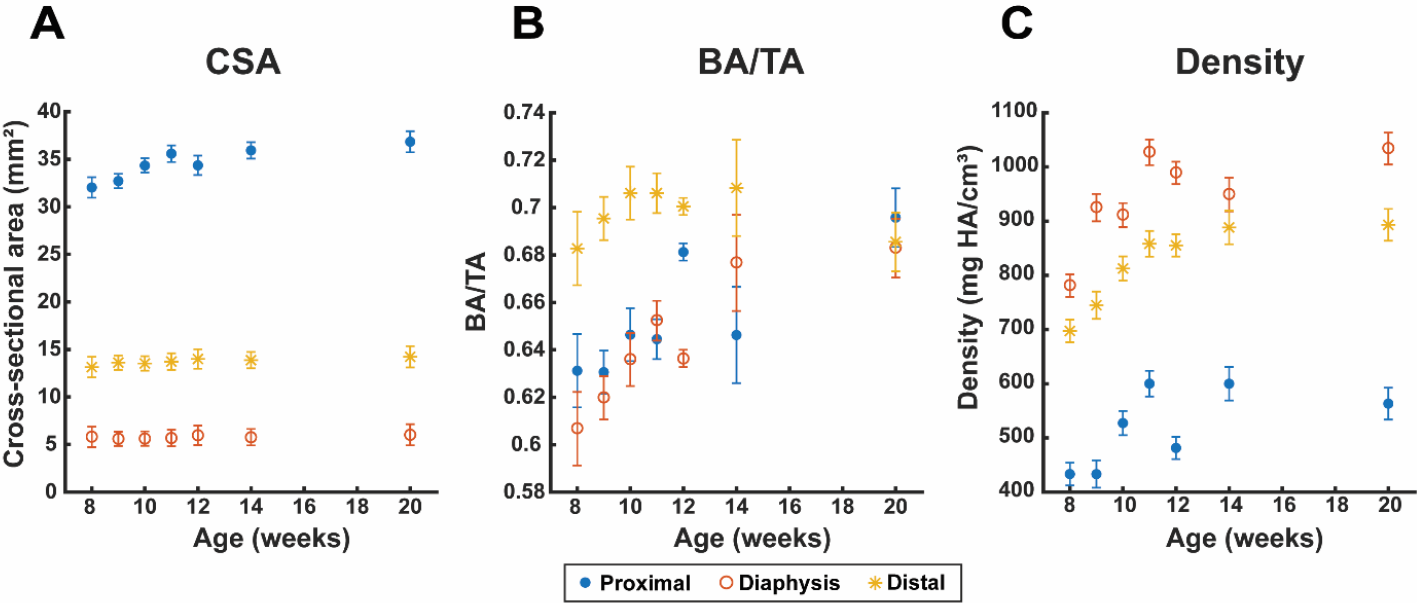
(A) Cross-sectional area, (B) bone area fraction, and (C) apparent density of proximal, middle, and distal regions were compared over time.

Body mass increased over time with a relatively constant slope of 6.13 ± 0.81 g/week. During level walking, the overall shape of the ground reaction force (GRF) was consistent during growth (Fig. 3A). At 8 weeks, the peak GRF was 0.69 ± 0.05 BW and increased by 21% at 12 weeks of age and 29% by 20 weeks of age (p<0.05) (Fig. 3B). The rate of increase was greatest between 8 and 12 weeks, after which the peak GRF began to plateau. The kinematics exhibited the greatest change between 8 and 10 weeks of age with the hip, knee and ankle joint angles becoming more flexed (p<0.05) (Fig. 3C,D). While the knee and ankle joint angles remained constant beyond ten weeks of age, the hip joint angle increased (less flexion) until fourteen weeks of age after which it remained constant.

**Figure 3.**
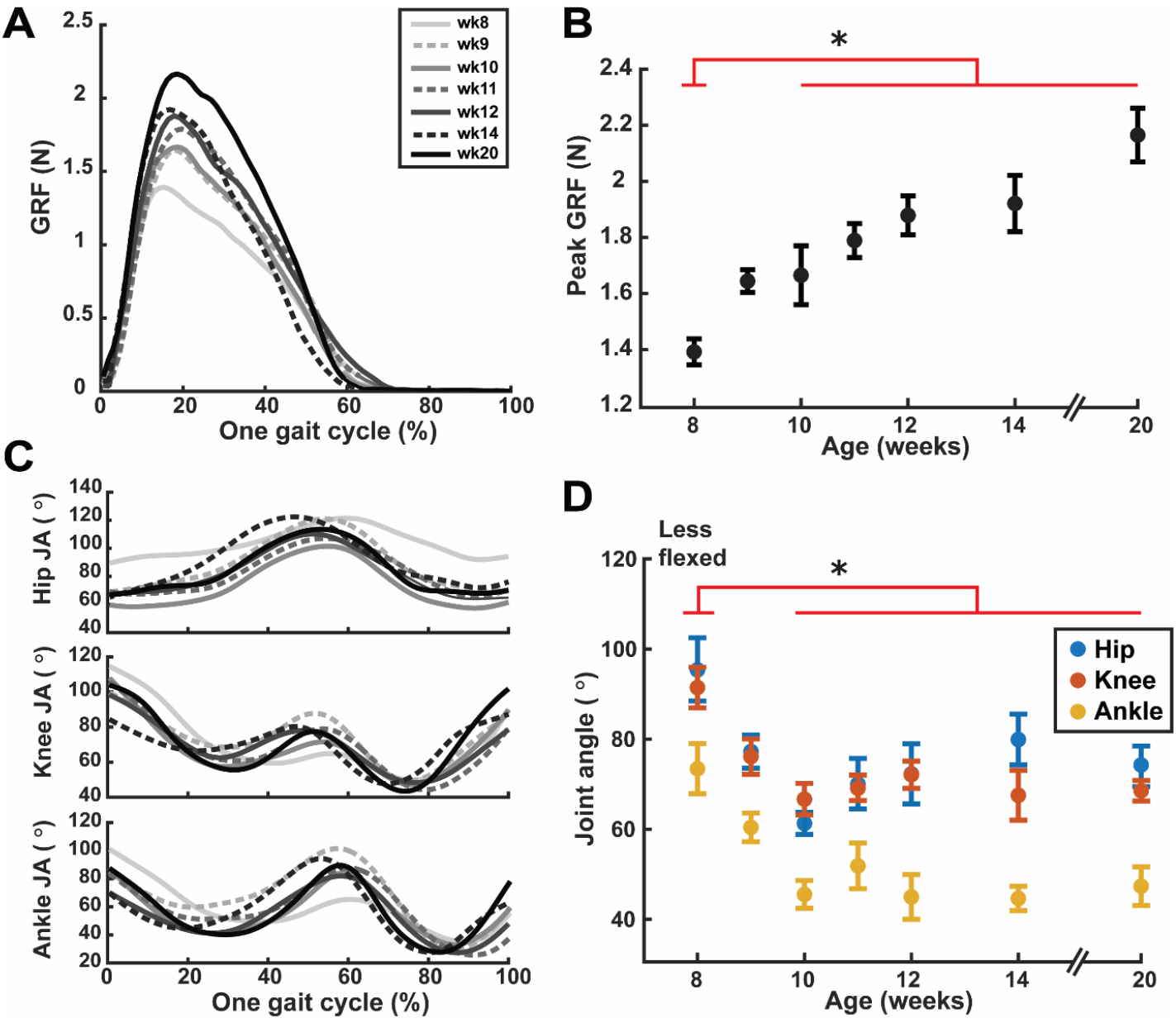
Kinematics and kinetics data of a representative rat. (A) Resultant ground reaction forces (GRF) of one gait cycle and (B) averaged peak GRF were measured during growth. (C) Three different joint angles during one gait cycle were measured at every time point and (D) joint angles at the time of the peak GRF were obtained at each time point and statistical analysis between time points was done. In the joint angle graph, 180 degree indicated fully extended joint and 0 degree indicated fully flexed joint.

Several bone morphological and density parameters were significantly correlated with gait parameters (Table 1). First, GRF was correlated with all parameters with the exception of distal BA/TA (r-range = 0.41-0.68, p<0.05). Kinematics tended to be more correlated to density rather than bone structure. Specifically, knee angle at the time of peak GRF was most correlated to density in all regions and proximal CSA (r-range = 0.42-0.62, p<0.05), followed by the ankle angle (r-range = 0.39-0.61, p<0.05). The hip angle was the least correlated to tibial properties (r-range = 0.41-0.46, p<0.05).

**Table 1.**
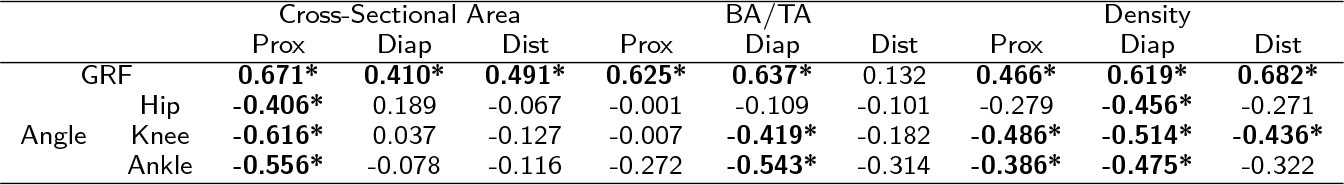
A statistical analysis to estimate the correlation coefficients between gait parameters and bone properties (* indicates p¡0.05).

The resistance of the tibia to compressive and bending loads was estimated using bone area and second moment of area, respectively. The proximal and diaphyseal bone areas increased with mass to the power of 0.74 and 0.78, which are higher than the isometric scaling but still include isometry within the 95% confidence interval (Fig. 4A). In the distal region, the scaling exponent of 0.48 for bone area was below the isometric scaling exponent. The second moment of area at the diaphysis had a scaling exponent of 1.81 which is close to the constant stress-based scaling exponent of 1.83 and the 95% confidence interval excluded the isometric expectation. The proximal and distal regions of the tibia had lower scaling exponents (1.1 and 1.08, respectively) than the isometric scaling exponent of 1.33 (Fig. 4B).

**Figure 4.**
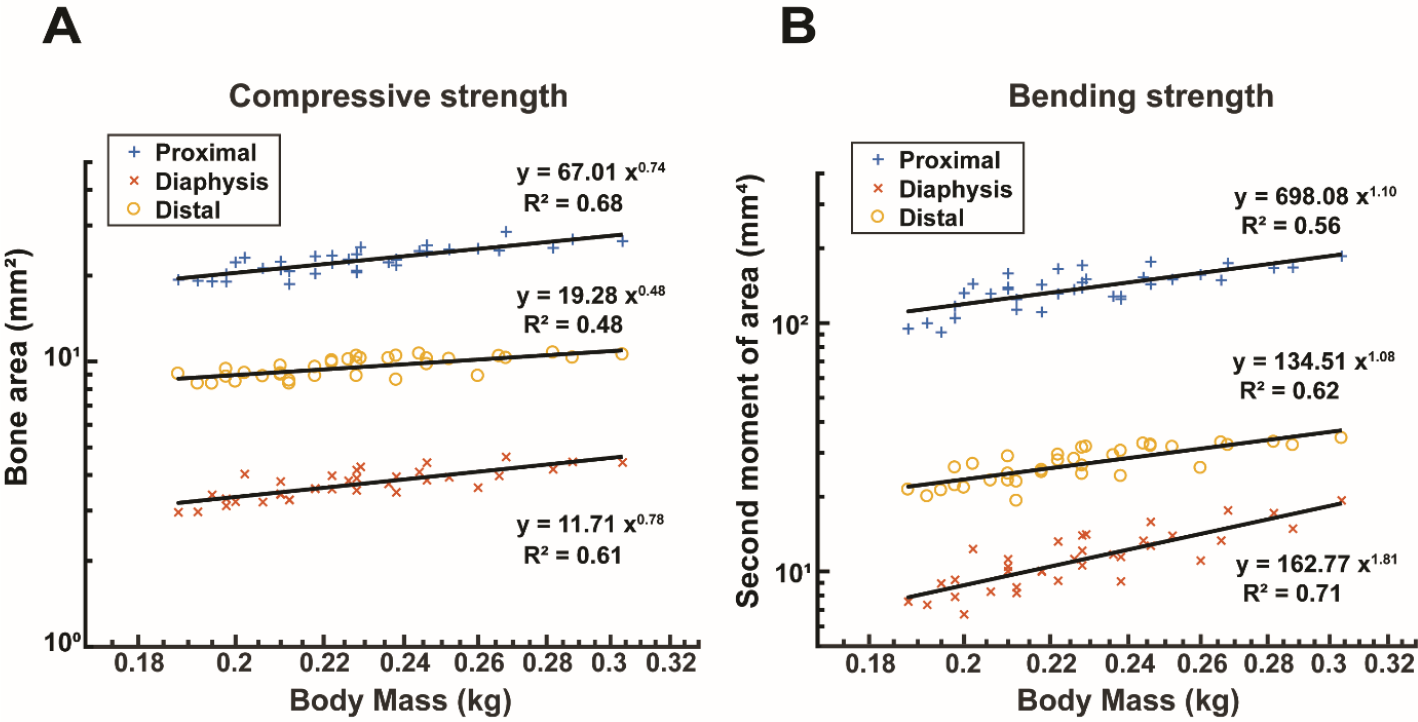
Power regression in each (A) cross-sectional bone area and (B) second moment of area with respect to body mass. Both parameters were fitted with a power law of the form, *y* = *ax^b^*.

The finite-element based analyses of bending and compressive strength revealed similar results as the scaling analyses. Overall, the mean strain energy density in the diaphysis decreased under bending and was variable under compressive loading. When evaluated beyond 10 weeks of age (when posture, total CSA, and density was constant) the mean strain energy density increased by 16% under compressive loads (*R*^2^ = 0.958) but decreased by 10% under bending loads (*R*^2^ = 0.998) (Fig. 5).

**Figure 5.**
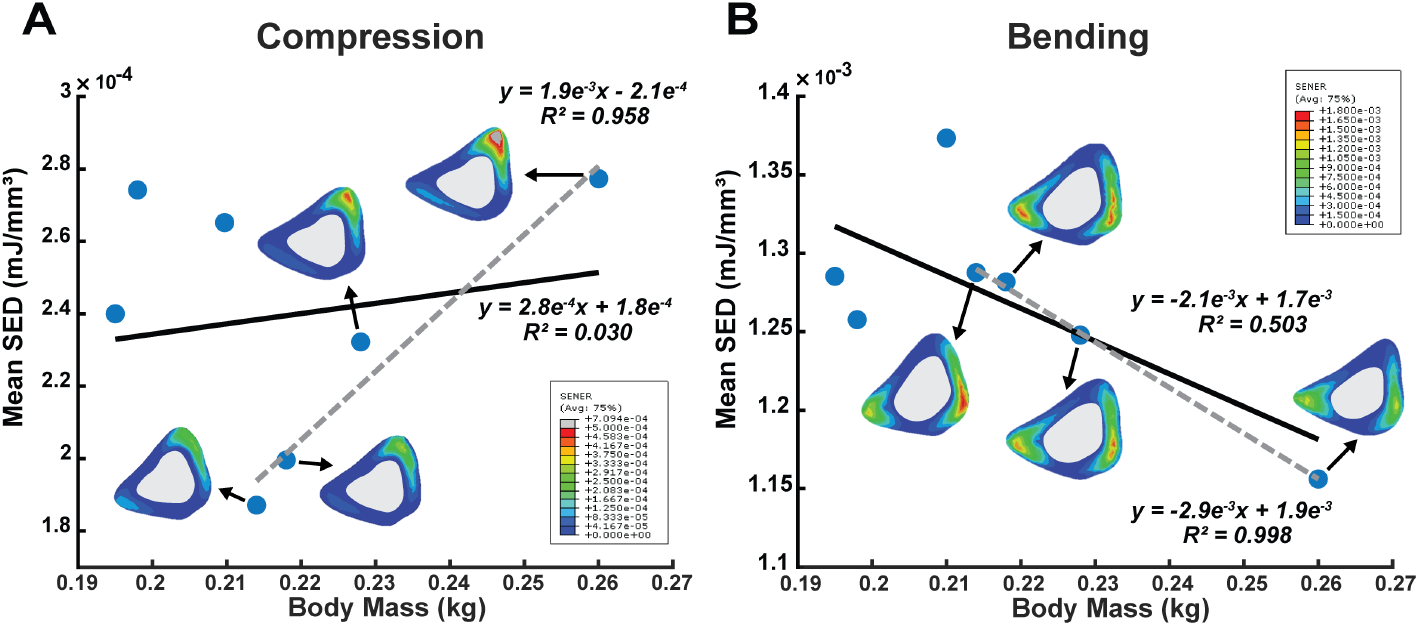
Mean strain energy density at tibial midshaft under (A) compression and (B) four-point bending.

## Discussion

During the relatively short period of growth, body size increases substantially providing an opportune window to understand how bone structure and material changes occur in response to (or in tandem with) increased loading. From our data, most significant changes in bone structure and density occurred early between 8 and 12 weeks of age, followed by less rapid or even no further changes in bone properties. This early growth phase was only evident in terms of size within the proximal CSA while the diaphysis and distal sizes remained relatively constant. However, changes in the distribution of bone (bone area fraction) was more consistent in all regions with clear increases in bone area fraction during early growth. Density increases were also most notable during early growth but was inversely related to size. The large proximal tibia had the lowest apparent density while the smaller diaphysis had the highest mineral density, suggesting that either an increase in bone fraction or density is required to accommodate increased loads but not both. However, the lower apparent density of the proximal tibia may be influenced by the partial volume effect associated with quantifying the density of trabecular bone and the relatively thinner cortex within this region.

The timing of structural and density changes during early growth was similar to the timing of changes in joint posture. Between eight and ten weeks, joint angles became more flexed after which posture was consistent. The bone properties were closely related to the knee and ankle joint angles and GRF during growth, which suggests that bone (re)modeling is responsive to changes in gait strategy in a focally specific manner. Joint angles were correlated to proximal and midshaft bone properties, but not distal properties. Changes in proximal CSA and bone area fraction continued one week beyond our observed postural changes, and may represent the time bone needs to undergo bone (re)modeling in response to the change in posture.

The convergence of bone area fraction by twenty weeks of age throughout the tibia suggests an attempt to evenly distribute bone material. At 20 weeks (93% of skeletal maturity) rats are almost fully grown, and our data suggest that gait posture has reached steady state. Therefore, beyond ten weeks when knee and ankle angles were constant, the orientation of loads on the tibia were likely consistent but only increasing in magnitude as reflected by the continual increase in ground reaction force. A body mass dependent increase of the peak GRF during growth was observed (Fig. 2), which is consistent with previous observations for many animals [33, 34].

The scaling and finite-element analyses suggest that the structural and material properties of the tibial diaphysis are organized to best accommodate bending loads. The second moment of area at the diaphysis followed the allometric scaling and the 95% confidence interval excluded the isometric expectation, while the scaling of bone area included the isometric exponent. This scaling analysis implies that rodent bone is (re)modeled in order to sustain the bending at the midshaft during growth rather than compressive loads, which was confirmed by the finite-element model results. Importantly, the FE model accounts for whole bone loading and a heterogeneous density distribution and our results imply that strain energy density is not constant in the diaphysis, but rather decreasing suggesting that the tibia is actively adapting to become stronger under bending but not in compression. Interestingly, only when the joint angles reached steady state, did the strain energy density show this relationship most clearly. The scaling analysis results are in general agreement with several previous studies of exercise induced strain distribution during growth. In vivo strain at chicken tibiotarsus and mandible of two different species of papio-nine primates was measured and the results of these studies supported that bones (re)modeled to maintain a static strain distribution during growth, corresponding with our results of stress similarity based scaling [37, 38]. There were some studies of negative allometric scaling, but they still demonstrated that long bones are being loaded primarily in bending through ontogeny [39, 40].

Throughout growth, the diaphyseal density remained the highest compared to the proximal and distal regions, but was relatively consistent in the second half of growth when posture was constant and mass was still increasing. Instead, the increase in bone loading due to mass was most clearly correlated to changes in bone area fraction in the diaphyseal region suggesting a stronger link between increasing mass and morphological changes. Increasing the amount of bone present, rather than increasing mineralization, may be the biologically efficient means of accommodating bending loads. Femoral and tibial midshafts have been shown to have the highest principal strains during walking or running in humans [41, 42], which supports the notion that the diaphysis requires allometric growth to minimize fracture risk. However, the degree to which bone is programmed at birth to be focally more adaptive based on genetics remains to be shown.

Importantly, rats and mice are the most commonly used animal models for the assessment of bone health, and one area of interest is in evaluating the bone response to mechanical loading for the treatment of bone disease or evaluation of pharmaceutical interventions in tandem with mechanical loading as a surrogate for exercise [43–46]. Exercise when young has been shown to be effective for inducing an adaptive response in bone [46], and our data has identified a time window in murine bone during which postural changes are associated with significant changes in bone properties. Several studies evaluating the effect of mechanical loading on bone use net axial compressive mechanical loads but often require increased strains or strain rates that are beyond those expected physiologically [43, 47–51]. However, our data suggest that the flexed postures at the tibia results in bending loads rather than compressive loads and may explain why a compressive loading model alone under physiological loading conditions does not result in an adaptive response.

There are several limitations to this study. Although the changes in gait strategy could be moderately correlated with bone properties change during growth, the gait kinematics and kinetics measured in this study do not necessarily reflect movements of rat normally. In this study, the rats spent most of their time in the cage, which was limited in space, even though walking is a general movement of murinae in cage [52]. Our analysis of bone area fraction and density is limited by the resolution of our CT data which may also include noise due to motion artifacts associated with in vivo imaging. The loading conditions used within the finite-element models are not reflective of in vivo loading. While we used animal and time specific data to provide relevant boundary conditions, they are not indicative of the direct loading of bone via muscle and joint forces during gait. The role of muscle forces in changes in bone properties during growth remains to be understood.

Positive allometry was observed in the second moment of area for diaphysis of rodent tibia and bone properties were correlated with gait strategy during growth. Scaling analyses used in this study are a powerful and useful tool in biomechanical modeling for understanding changes in bone properties with increasing body size and understanding how bone accommodates increases in mass.

## Competing interests

The authors declare that they have no competing interests.

## Acknowledgements

We are grateful for the assistance of Iwona Dobrucka, Wawrzyniec Dobrucki, Christian Konopka, Than Huynh, and Travis Ross in data collection and processing.

